# Systematic analysis of cilia characteristics and Hedgehog signaling in five immortal cell lines

**DOI:** 10.1101/2022.03.22.485292

**Authors:** Arianna Gómez, Julie Craft Van De Weghe, Malaney Finn, Dan Doherty

**Author notes:** Corresponding author (DD).

## Abstract

Dysfunction of the primary cilium, a microtubule-based signaling organelle, leads to genetic conditions called ciliopathies. Hedgehog (Hh) signaling is mediated by the primary cilium in vertebrates and is therefore implicated in ciliopathies; however, it is not clear which immortal cell lines are the most appropriate for modeling pathway response in human disease; therefore we systematically evaluated Hh in five commercially available, immortal mammalian cell lines: ARPE-19, HEK293T, hTERT RPE-1, NIH/3T3, and SH-SY5Y. All of the cell lines ciliated adequately for our subsequent experiments, except for SH-SY5Y which were excluded from further analysis. hTERT RPE-1 and NIH/3T3 cells relocalized Hh pathway components Smoothened (SMO) and GPR161 and upregulated Hh target genes in response to pathway stimulation. In contrast, pathway stimulation did not induce target gene expression in ARPE-19 and HEK293T cells, despite SMO and GPR161 relocalization. These data indicate that human hTERT RPE-1 cells and murine NIH/3T3 cells, but not ARPE-19 and HEK293T cells, are suitable for modeling the role of Hh signaling in ciliopathies.

## Background

Primary cilia are microtubule-based organelles that transduce light, mechanical, and chemical signals into cells (1). Protruding from the surface of most cell types, primary cilia mediate multiple signaling pathways, including Hedgehog (Hh), that play important roles in development and tissue homeostasis. Cilium dysfunction results in conditions called ciliopathies, and while abnormal Hh signaling has been described in models of ciliopathy-associated genes, details of the disease mechanism have not been clearly elucidated. (2). Immortal cell lines are frequently used to study primary cilium biology and the mechanisms underlying ciliopathies (3–6); however, the strengths and weaknesses of commonly used human cell lines have not been systematically compared. The goal of this study was to identify immortal cell lines suitable for studying Hh signaling in human health and disease.

In vertebrates, the primary cilium modulates the canonical Hh signaling pathway through the coordinated translocation of Hh pathway proteins in and out of the cilium (reviewed in (7)). In the absence of Hh ligand, ciliary localization of the Patched (PTCH) receptor indirectly represses ciliary localization of Smoothened (SMO), maintaining the pathway in an inactive state. This repression is reinforced by ciliary GPR161, an orphan constitutively active G-coupled protein receptor and negative regulator of the pathway, which maintains high ciliary cyclic AMP (cAMP) levels which in turn increases protein kinase A (PKA) activity. Increased PKA activity lead to proteolytic cleavage of GLI transcription factors into their repressor forms, maintaining expression of target genes low (8). During pathway stimulation, Hh ligand binds PTCH, promoting its export from the cilium, which results in SMO entry and GPR161 exit. In concert, GLI2/3, KIF7, and Suppressor of Fused (SUFU) localization to the ciliary tip increases, and is associated with reduced GLI2/3 cleavage (9,10). Uncleaved GLI2/3 activator forms translocate to the nucleus and induce transcriptional targets including *GLI1* and *PTCH1* (11,12). In this work, we focus on canonical Hh signaling because it has been implicated in the mechanisms underlying ciliopathies across multiple model systems. We compare two steps in the Hh pathway in immortal cell lines: initial ciliary localization of Hh pathway components and downstream Hh target gene expression.

## Materials and Methods

### Cell culture

Cell lines were obtained from the American Type Cell Collection (ATCC, www.atcc.org) and grown in cell specific medium. ARPE-19 (ATCC, CRL-2302) and hTERT RPE-1 (ATCC, CRL-4000) cells were cultured in DMEM/F12 with 10% fetal bovine serum (FBS) and 1% penicillin-streptomycin. HEK293T (ATCC, CRL-3216) cells were cultured in DMEM with 10% FBS and 1% penicillin-streptomycin. NIH/3T3 (ATCC, CRL-1658) cells were cultured in DMEM with 10% bovine calf serum and 1% penicillin-streptomycin. SH-SY5Y (ATCC, CRL-2266) cells were cultured in 1:1 Eagle’s Minimum Essential Medium:F12 with 10% FBS and 1% penicillin-streptomycin. 0.05% trypsin was used for all cell dissociation.

### Time course assay

For each set of experiments, cells were seeded on pairs of coverslips coated with 0.3mg/mL poly-D-lysine and allowed to grow for 2-3 days, until they reached 60-80% confluency. We define batches as cells that were grown, treated, and collected at the same time. After reaching the desired confluency, we serum starved the first pair of coverslips that would be starved for the longest time point (96-or 72-hours). Serum starvation included removing the complete growth medium, rinsing with PBS, and adding serum free medium. This step was repeated for each time point, with all coverslips from the same cell line and batch collected at the end of the time course (0 hour serum starvation). We placed the plates holding the coverslips on ice for 10 minutes, then fixed coverslips using 4% paraformaldehyde for 5 minutes followed by permeabilization using cold methanol for 3 minutes and stored in PBS at 4°C until ready to stain. We only stained one coverslip from the pair, leaving the replicate available as a backup.

For HEK293T cells, we incubated the 0.3 mg/mL poly-D-lysine coated coverslips with 0.05% gelatin for 15-20 minutes prior to seeding cells to promote adhesion since this cell line is semi-adherent. Otherwise, we followed the same seeding and fixation method described above.

All plots were generated using Plots of Data and Super Plots of Data (13,14)

### Hedgehog pathway protein localization

Cells in the same batch were seeded on pairs of coverslips and allowed to expand until they reached 60-80% confluency, as above. Cells were serum starved for a total of 48 hours. After the initial 24 hours of serum starvation, we replaced the media on half of the coverslips with serum free medium + 1μM Smoothened Agonist (SAG) (Millipore, 566661). We placed the plates holding the coverslips on ice for 10 minutes, then fixed coverslips using 4% paraformaldehyde for 5 minutes followed by permeabilization using cold methanol for 3 minutes and stored in PBS at 4°C until ready to stain. We only stained one coverslip from the pair, leaving the replicate available as a backup.

### Quantitative immunofluorescence

We blocked coverslips in 2% bovine albumin serum (BSA) in phosphate buffered solution (PBS) for 20 minutes at room temperature. Time course assay coverslips were incubated with mouse anti-acetylated α-tubulin (1:2000, Sigma Aldrich, T6793) and rabbit anti-ARL13B (1:800, Proteintech, 17711-1-AP) antibodies diluted in blocking buffer. Hedgehog pathway protein localization coverslips were incubated with either 1) mouse anti-SMO (1:50, Santa Cruz Biotechnology, sc-166685) and rabbit anti-ARL13B (1:800, Proteintech, 17711-1-AP) antibodies or 2) mouse anti-ARL13B (1:200, NeuroMab, 75-287) and rabbit anti-GPR161 (1:200, Proteintech, 13398-1-AP) antibodies diluted in blocking buffer. Coverslips were incubated with primary antibodies for 1 hour at room temperature or at 4°C overnight, then washed in PBS for 5 minutes, three times. Coverslips were incubated with the following secondary antibodies at 1:400 dilution for 1 hour at room temperature: Goat anti-Rabbit IgG, Alexa Fluor 488 (Thermo Fisher Scientific #A11008) and Donkey anti-Mouse IgG, Alexa Fluor 568 (Thermo Fisher Scientific, #A10037). After incubation, we washed coverslips in PBS for 5 minutes, three times. Coverslips were mounted on slides using a Fluoromount with DAPI (Invitrogen, #00-4959-52), then sealed with nail polish after sitting for at least 1 hour.

We quantified ciliary protein localization using a validated protocol previously established in the laboratory (15). We imaged coverslips using the same microscopy settings to acquire z-stack images for >20 cells with cilia for each condition and batch using identical microscope settings. We converted z-stack images to sum-projections and randomized these images using the FIJI (16) script, Filename_randomizer (https://imagej.nih.gov/ij/macros/Filename_Randomizer.txt) to minimize bias between cell lines and conditions. Prior to data collection, we checked a subset of images to ensure that the cilium marker (ARL13B) and protein of interest (SMO or GPR161) signals were adequate for measurement. Blinded to the condition, we drew a cilium mask in the cilium marker channel (ARL13B) using summed Z-stack images in ImageJ. We used this mask to measure the signal intensity for the protein of interest and measured the signal intensity of an adjacent region to subtract background signal. Normalized fluorescence intensity was calculated using the following formula:

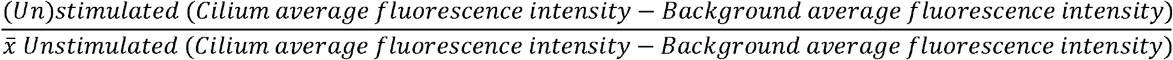

Finally, we unblinded the data and visually inspected the images to ensure that our qualitative assessment of ciliary signal was consistent with the quantitative data. We also used the same data set to determine proportion of cells with cilia and cilium length in baseline and stimulated cells.

### *GLI1* and *PTCH1* qPCR

Cells were grown in pairs of T-75 cell culture flasks until 60-80% confluent, then starved for a total of 48 hours. After the initial 24 hours, half the flasks had their media replaced with serum free medium + 1µM SAG. We dissociated the cells and extracted RNA using the Aurum Total RNA mini kit (Biorad, 7326820). We measured RNA concentration using a spectrophotometer and only included RNA that had an A260/280 >1.8. cDNA was generated using the BioRad iScript cDNA Synthesis kit. We set up qPCR reactions using the PowerUp SYBR Green Master Mix (Thermo Fisher Scientific, #A25741). For cell lines that required the touchdown qPCR protocol, we followed the protocol outlined in Zhang et al. (17). Briefly, the touchdown qPCR protocol went as follows: 50C (2 min), 95C (2 min), 4x [95C (20 sec), 65C (10 sec, decrease 3C/cycle, 72C (1 min)], 40x [95C (15 sec), 55C (15 sec), 72C (1 min)], ending with a melt temperature curve. qPCR data acquisition was performed on the Bio-Rad CFX96 Touch Real-Time PCR Detection System.

Human *GLI1* primers

Forward 5’-GATGACCCCACCACCAATCAGTAG-3’

Reverse 5’-AGACAGTCCTTCTGTCCCCACA-3’

Human *PTCH1 primers*

Forward 5’-GAGCACTTCAAGGGGTACGA-3’

Reverse 5’-GGAAAGCACCTTTTGAGTGG-3’

Mouse *Gli1* primers

Forward 5’-CCGACGGAGGTCTCTTTGTC-3’

Reverse 5’-GCGTCTCAGGGAAGGATGAG-3’

Human *Ptch1 primers*

Forward 5’-GAGCAGATTTCCAAGGGGAAG -3’

Reverse 5’-CCACAACCAAAAACTTGCCG -3’

### Reference gene identification

To ensure robust data acquisition, we identified an ideal reference gene for each cell line. We identified 10 human candidate reference genes and 9 mouse candidate reference genes (Table 1) to evaluate expression stability between baseline and stimulated cDNA. To select a reference gene, we evaluated the cycle threshold (CT) of three starting primer concentrations, and cDNA dilution curve efficiency (18).

**Table 1.**
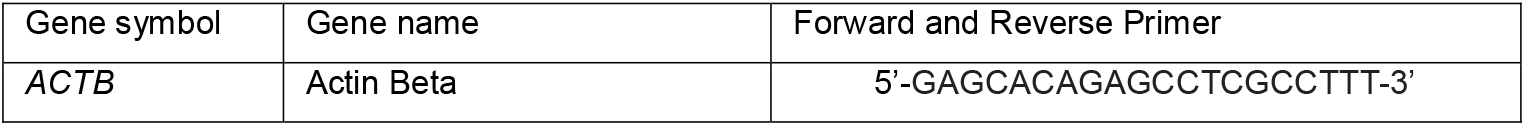
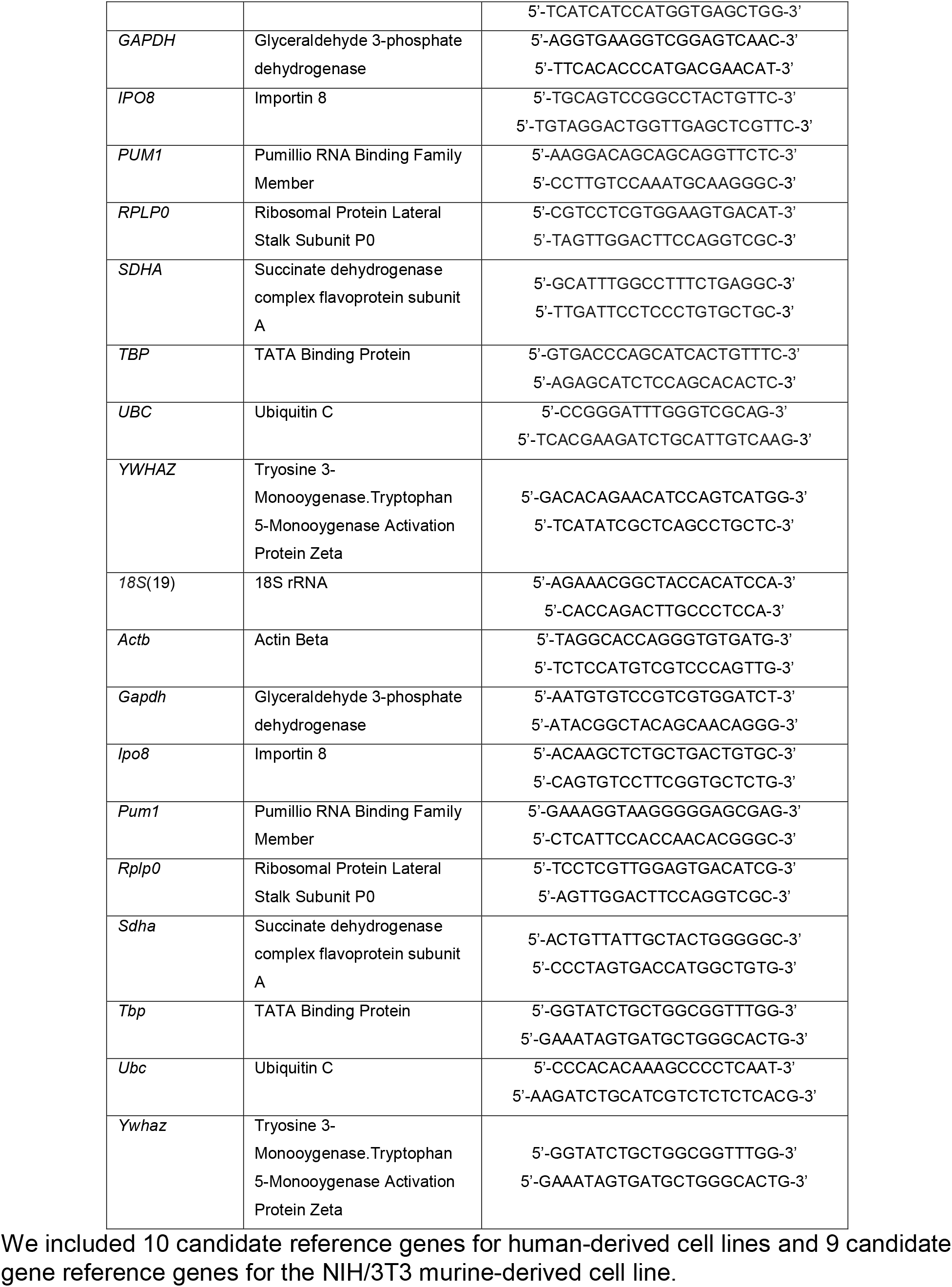
Candidate qPCR reference gene names and primer sequences.

In our initial screen, we were looking for 1) small ΔCT between unstimulated and stimulated within each cell line, indicating unchanged expression level between experimental conditions, and 2) a CT between 20-30 cycles so that expression levels were similar to *GLI1* and *PTCH* expression. We optimized primer concentration by diluting forward and reverse primers sets to three concentrations, 3µM, 5µM, and 7µM. Next, we determined primer efficiency by performing a cDNA dilution series. We followed widely accepted primer efficiency standards, which includes an efficiency of 80-120% and R^2^≥ 0.99.

During assay validation, we determined that *GLI1* in ARPE-19, HEK293T, and NIH/3T3 cells amplified at >29 cycles, close to the limit of reliable detection; therefore, we performed a modified qPCR assay in these cell lines, which included a touchdown step prior to the 40-cycle amplification. This lowered the detection level by 3 cycles in ARPE-19 and HEK293T cells. In NIH/3T3 cells, we did not see a difference in unstimulated cycle values and only saw a difference of 1 cycle in stimulated cells.

### Statistical analysis

An unpaired two-tailed Student’s *t-*test was performed for comparison between unstimulated and stimulated cells with a hypothesized mean difference of 0. α level was set at 0.05. *P* < 0.05 was significant. Symbols for significance represent *p*-values from 0.01 to 0.05 (*), 0.001 to 0.01 (**), and <0.001 (***).

## Results

### Cell line selection

We identified candidate models for human Hh signaling by evaluating the following characteristics in commercially available cell lines: human-derived, evidence of ciliation, immortal, required imaging characteristics (adherent, monolayer growth), evidence of Hh pathway response, and genetic tractability. ARPE-19, HEK293T, hTERT RPE-1, NIH/3T3, and SH-SY5Y cells met most criteria (Table 2). We included murine NIH/3T3 cells as a non-human control since these cells are frequently used to model Hh pathway function (6,20–22).

**Table 2.** Desired cell line characteristics for modeling Hh signaling.

|  | ARPE-19 | HEK293T | hTERT<br>RPE-1 | SH-SY5Y | NIH/3T3 |
| --- | --- | --- | --- | --- | --- |
| Cell origin | Retinal pigment epithelial cells | Embryonic kidney cells | Retinal pigment epithelial cells | Bone marrow derived Neuroblastoma cells | Murine embryonic fibroblast cells |
| Human-derived | Y | Y | Y | Y | N |
| Ciliated | Y | Y | Y | Y | Y |
| Immortal | Y | Y | Y | Y | Y |
| Adherent | Y | S | Y | Y | Y |
| Monolayer growth | Y | N | Y | N | Y |
| Hh pathway response | Y | Y | Y | Y | Y |
| Gross chromosomal abnormalities | N | Y | N | N | N |
| Genetic tractability | Y | Y | Y | NR | Y |

We searched the literature for evidence of the desired characteristics listed. ATCC often provided information about the following categories: human-derived, immortal, gross chromosomal abnormalities. When considering any known response to Hedgehog signaling, this included assays that measured response differently from the assays discussed in this paper (i.e. GLI3 processing, GLI luciferase assays). Y= Yes, N= No, S = semi-adherent, NR= Not reported.

### Proportion of cells with cilia

While all the cell lines have been reported to ciliate, the time course of ciliation for different cell lines is scattered across studies (23,24). Therefore, we aimed to define the optimal serum starvation conditions for Hh signaling experiments by determining the proportion of ciliated cells across time points after serum starvation: 0, 8, 16, 24, 48, 72, and 96 hours. Serum starvation removes growth factors available to cells, causing cells to drop out of the cell cycle and promoting ciliogenesis. For Hh imaging assays, we wanted cells to be in a monolayer layer, but we were unable to measure the 96-hour time point for HEK293T cells, because they became overgrown when grown for >72 hours under standard conditions.

Baseline ciliation varied between cell types, ranging from ∼5% in SH-SY5Y to ∼62% in hTERT RPE-1 (Fig 1). Serum starvation was associated with >15% higher ciliation rates over baseline in ARPE-19, hTERT RPE-1, and NIH/3T3 cell lines. Peak ciliation in these cell lines typically occurred 48 hours post-starvation and ranged from 56% in ARPE-19 to 97% in NIH/3T3. In contrast, HEK293T and SH-SY5Y ciliation rates were the same with and without serum starvation.

**Fig 1.**
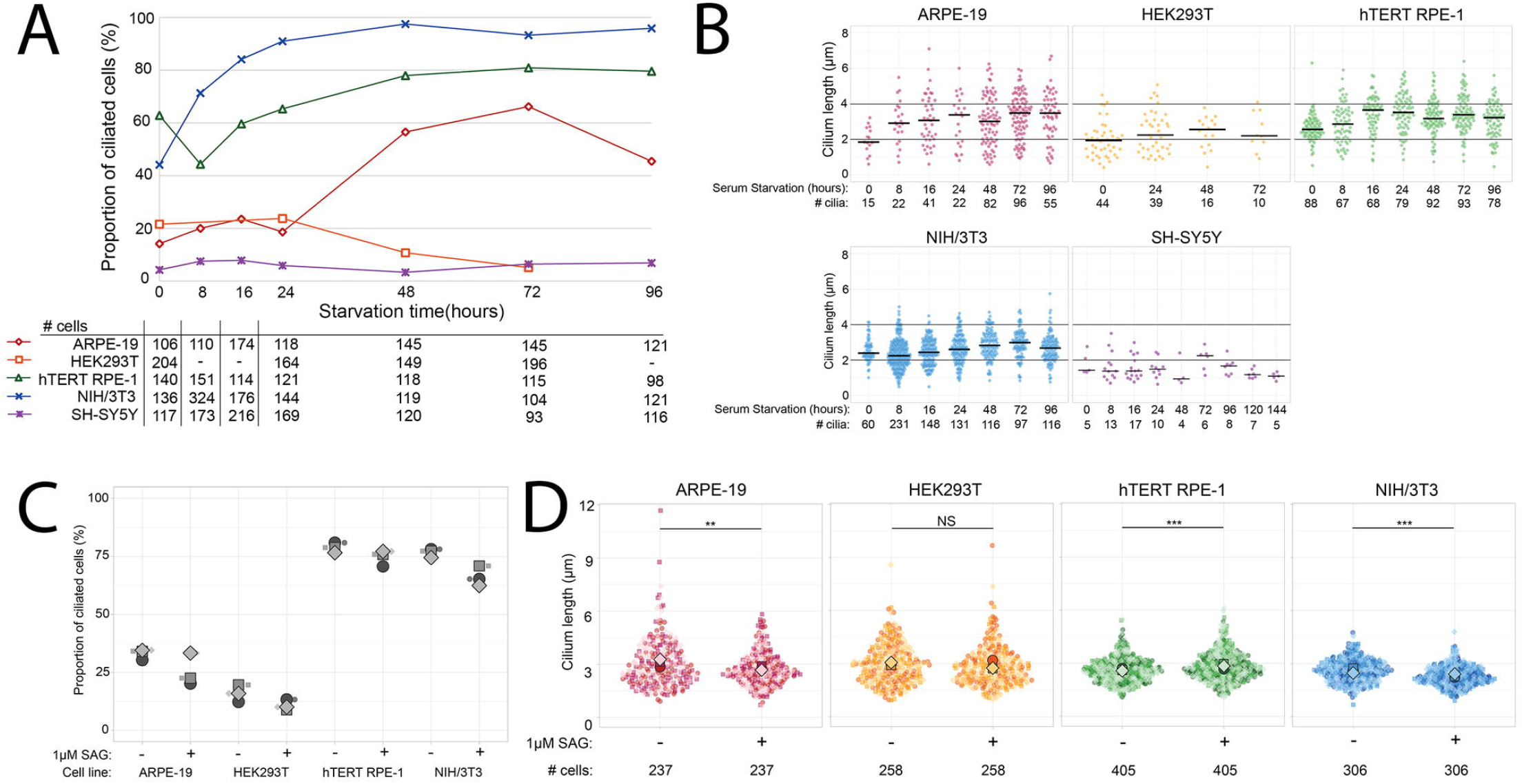
Ciliation time course and cilia length for immortal cell lines. A) Proportion of ciliated cells at 0, 8, 16, 24, 48, 72, and 96 hours post serum starvation. The numbers of cells counted at each time point are listed below each column of the graphs. B) Cilium length across time points. Black bars across graphs delineate 2 and 4μm. Black bars in each column represent median length. C) Proportion of ciliated cells without and with stimulation. Each symbol represents the proportion of ciliated cells in one batch. D) Cilium length without stimulation and with stimulation. Large symbols = median cilium length within a batch. Small symbols = cilium length of one cilium. One coverslip was imaged for each cell line and time point. The total number of cells counted across three batches are listed below each column of the graphs.

SH-SY5Y cells ciliated at low proportions through 96 hours of starvation, prompting us to evaluate 120- and 144-hour time points to determine whether ciliation was delayed compared to other cell lines. Even with prolonged starvation, <8% of SH-SY5Y cells had cilia at each time point (S1 Fig).

We and others have previously documented proportions of ciliated HEK239T cells >20% (25). To determine whether the low proportions of cells with cilia was a characteristic of the specific isolate of HEK293T cells, we repeated the time course experiment using three different isolates of HEK293T cells in our collection (S2A Fig). Baseline ciliation rates were similar to our initial experiment (∼20%) and increased to 32-39% by 24 hours. At 48 and 72 hours, the proportion of cells with cilia was lower than at 24 hours. We compared images of areas monolayer and multilayer growth, finding that more densely grown areas had more cilia (S2B, S2C Fig). For the Hh experiments described below, we continued with monolayer cultures to allow for accurate quantitative immunofluorescence measurements.

### Cilium length

We measured cilium length across all timepoints (Fig 1B). Cilium length was shorter at 0 hours for ARPE-19, HEK293T, hTERT RPE-1, and NIH/3T3 cells, stabilizing between 8-24 hours, while cilium length was similar across timepoints in SH-SY5Y cells. Maximum median length for each cell line was 3.0μm for ARPE-19, 2.1μm for HEK293T, 3.2μm for hTERT RPE-1, 2.8μm for NIH/3T3, and 1.4μm for SH-SY5Y cells. Based on the low ciliation rate and exceptionally short cilia, we did not continue evaluating SH-SY5Y cells for Hh pathway response.

### Hedgehog pathway response

We next measured canonical Hh pathway activity in the cell lines using two Hh pathway assays commonly reported in the literature: 1) upstream localization of Hh pathway effectors SMO and GPR161 into and out of the cilium, respectively (8,26), and 2) downstream target gene (*GLI1* and *PTCH1)* induction (11).

#### Hh pathway protein localization

We measured the ciliary localization of SMO and GPR161 in unstimulated and Smoothened agonist (SAG) stimulated cells using our established quantitative immunofluorescence (qIF) protocol that determines relative protein localization between different cell lines and conditions (15).

#### Proportion of cells with cilia and cilium length in response to Hh stimulation

Proportion of ciliated cells and cilium length were moderately affected by pathway stimulation (Fig 1C,1D). Proportion of ciliated cell was lower in stimulated cells compared to unstimulated cells across cell lines in a majority of batches. Cilium length was significantly decreased in ARPE-19 and NIH/3T3 cells, significantly increased in hTERT RPE-1 cells, and was not significantly different in HEK293T cells.

#### SMO localization

Normalized mean ciliary SMO signal was significantly higher in stimulated cells compared to unstimulated cells across all cell lines, with fluorescence intensity reported in arbitrary units (au): 1.5 – 2.6 au in ARPE-19, 1.2 – 2.7 au in HEK293T, 2.1 – 2.9 au in hTERT RPE-1, and 7.0 – 10.3 au in NIH/3T3 cells (Fig 2A). In ARPE-19 and HEK293T cells, SMO ciliary fluorescence intensity was weaker and background signal was higher compared to hTERT RPE-1 and NIH/3T3 cells (S2A Fig).

**Fig 2.**
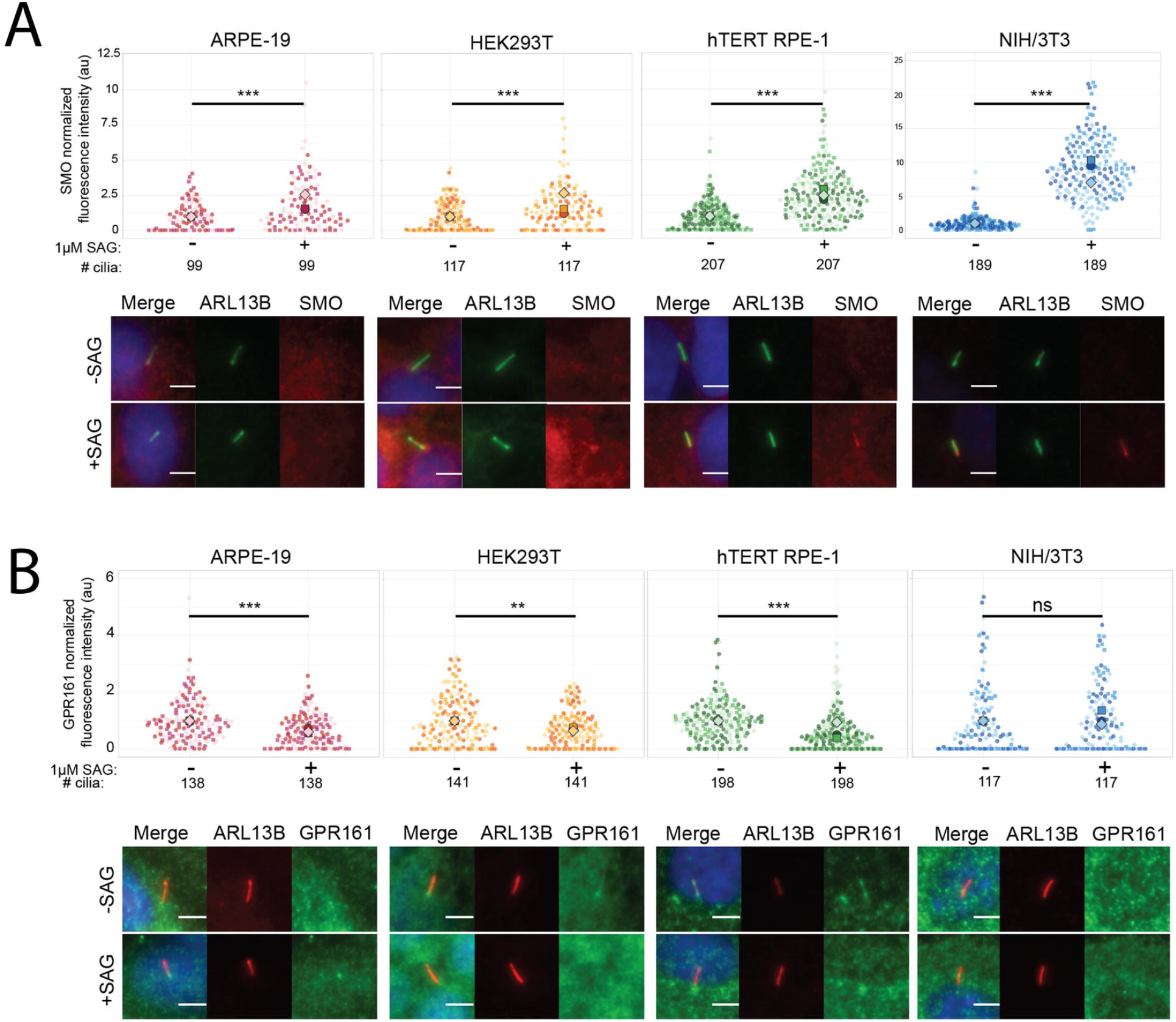
SMO and GPR161 localization in response to Hh pathway stimulation. Normalized fluorescence intensity (au) of A) SMO and B) GPR161 with and without pathway stimulation. Large symbol = median normalized fluorescence intensity for individual coverslip data. Small symbol = normalized fluorescence intensity of one cilium. Measurement from three separate batches are indicated by a unique symbol shape and color. Total number of cilia measured listed in the second row below the graph. Representative images are shown below graphs for unstimulated (-SAG) and stimulated (+SAG) cells. Two-tailed Student’s *t-*test was performed. *P*-values represented significant differences if they ranged from 0.01 to 0.05 (*), 0.001 to 0.01 (**), and <0.001 (***).

#### GPR161 localization

We detected cilium-specific GPR161 signal in HEK293T and hTERT RPE-1 cells, a faint cilium-specific signal in ARPE-19 cells, and no cilium-specific signal in NIH/3T3 cells (Fig 2B), so we could not measure GPR161 response in NIH/3T3 cells. Across three batches, normalized mean GPR161 intensity was lower with SAG stimulation: 0.57-0.73 au in ARPE-19, 0.64 -0.76 au in HEK293T, and 0.40 – 0.95 au in hTERT RPE-1 (Fig 2B). For one of the three hTERT RPE-1 passages (Batch 3), the GPR161 signal was the same with and without stimulation. We suspect that this was due to an undetermined technical issue since hTERT RPE-1 cells responded when we repeated the experiment on an additional passage (S2C Fig).

#### Hh target gene expression

*GLI1* and *PTCH1* are direct transcriptional outputs of the canonical Hh pathway and are commonly used to measure pathway activity (11). To measure *GLI1* and *PTCH1* expression, we optimized our quantitative PCR assays using the 2^-ΔΔ*;CT*^ method (27).

To accurately measure transcriptional responses to pathway stimulation using the 2^-ΔΔ*CT*^ quantitative PCR method, we identified suitable reference genes whose expression levels were similar to *GLI1* and *PTCH1* (>20 cycles, <30 cycles) and did not change with pathway stimulation in each cell line. We searched the literature for commonly used reference genes, identifying ten candidates for human cell lines and nine candidates for the mouse cell line (primers listed in Methods). We chose the genes with the most similar expression levels in stimulated and unstimulated cells, with the amplification cycles closest to *GLI1* and *PTCH1*: *PUM1* for ARPE-19 cells, *TBP* and *PMBS* for HEK293T cells, *TBP* for hTERT RPE-1 cells, and *Sdha* for NIH/3T3 cells (S4 Fig). We also used murine specific *Gli1* and *Ptch1* primer sets for NIH/3T3 cells. Although we tested *TBP* and *PMBS* for HEK293T, and used *PMBS* in our experiment, neither was ideal: *TBP* expression was variable in stimulated cells, and *PMBS* expression varied with stimulation in one out of the three batches (S4 Fig).

In hTERT RPE-1 cells, *GLI1* expression was 2.8 to 3.8-fold higher and *PTCH1* expression was 1.7 to 1.9-fold higher with stimulation (Fig 3). In contrast, stimulation was not associated with marked expression differences in ARPE-19 cells (*GLI1* 0.9 to 1.4-fold, *PTCH1* 1.0 to 1.1-fold) or HEK293T cells (*GLI1* 0.5 to 1.7-fold, *PTCH1* 0.3 to 0.7 fold), consistent with the lack of relocalization of SMO and GPR161 with SAG stimulation. Strikingly, *Gli1* expression was 1993 to 3034-fold higher with stimulation in NIH/3T3 cells. Similarly, *Ptch1* expression was also markedly higher (46 to 70-fold).

**Fig 3.**
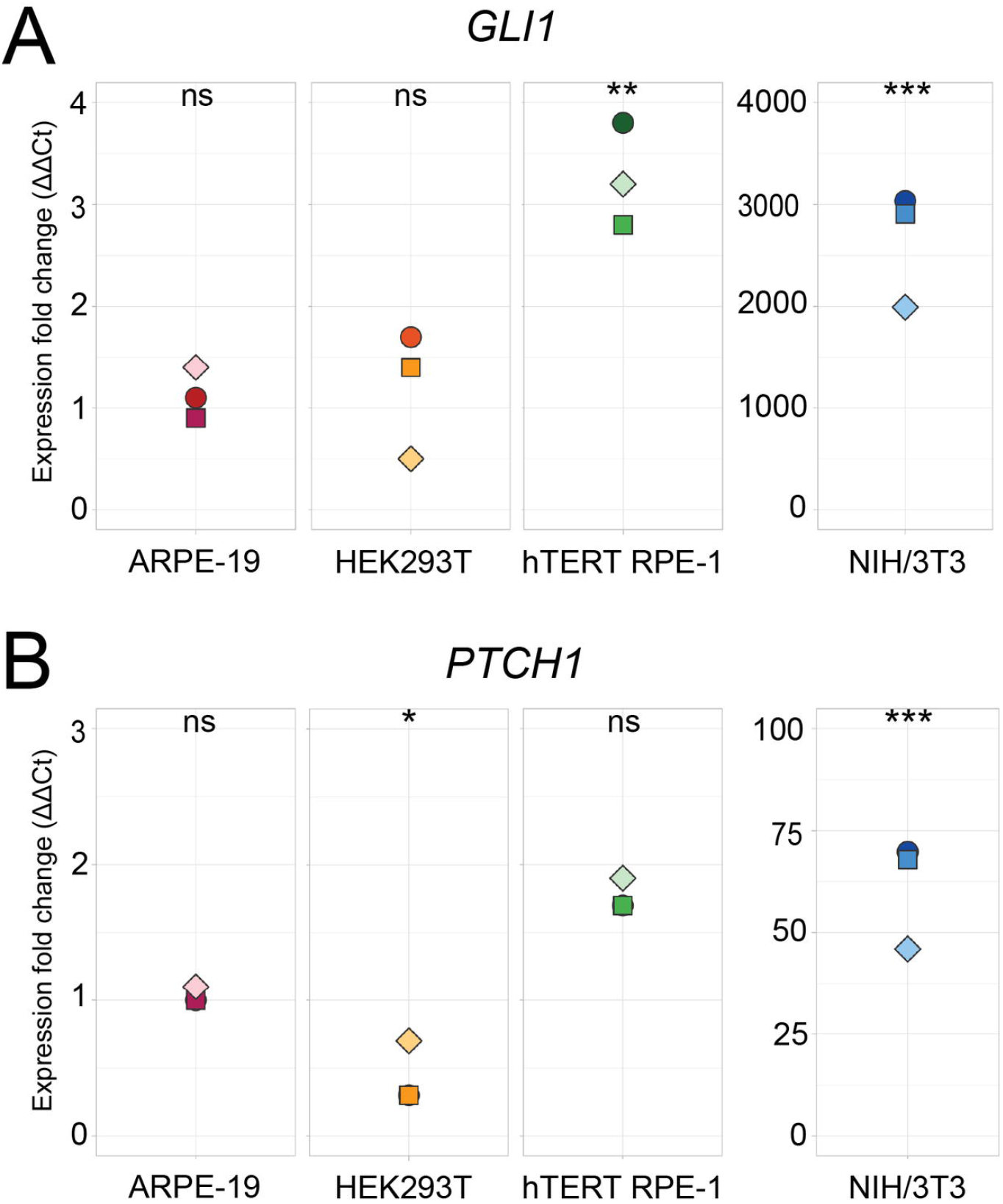
*GLI1* and *PTCH1* expression in response to Hh pathway stimulation. A) *GLI1* and B) *PTCH1* target gene induction. Each data point represents the expression fold change of stimulated cells from the same batch. Expression fold change was normalized to unstimulated values, which were set to 1. Two-tailed Student’s *t-*test was performed. *P*-values represented significant differences if they ranged from 0.01 to 0.05 (*), 0.001 to 0.01 (**), and <0.001 (***).

## Discussion

This systematic evaluation of cilium characteristics and Hh signaling in five commercially available, immortal, mammalian cell lines revealed that hTERT RPE-1 and NIH/3T3 cells are suitable for modeling Hh signaling (Table 3). By using the same assays in different cell types, we determined differences in response to Hh stimulation. Based on our experiments, SH-SY5Y cells were not suitable because of their low ciliation rate and very short cilia. ARPE-19 and HEK239T cells localizaed the Hh pathway proteins SMO and GPR161 in response to stimulation; however, they did not upregulate the Hh target genes *GLI1* and *PTCH1*. Only hTERT RPE-1 and NIH/3T3 cells ciliated well and showed robust responses to stimulation in both sets of assays. Our experiments also demonstrate that Hh pathway stimulation moderately impacts ciliation rates and cilium length across all of the cell lines, but not in a consistent manner across batches. We observed a range of responses to stimulation, from minimally responsive (ARPE-19 and HEK293T), to modestly responsive (hTERT RPE-1) to robustly responsive (NIH/3T3) (Table 3). It is not clear whether this represents species-specific or cell line-specific differences, but it does indicate that results should be validated across model systems before they can be considered generalizable.

**Table 3.**
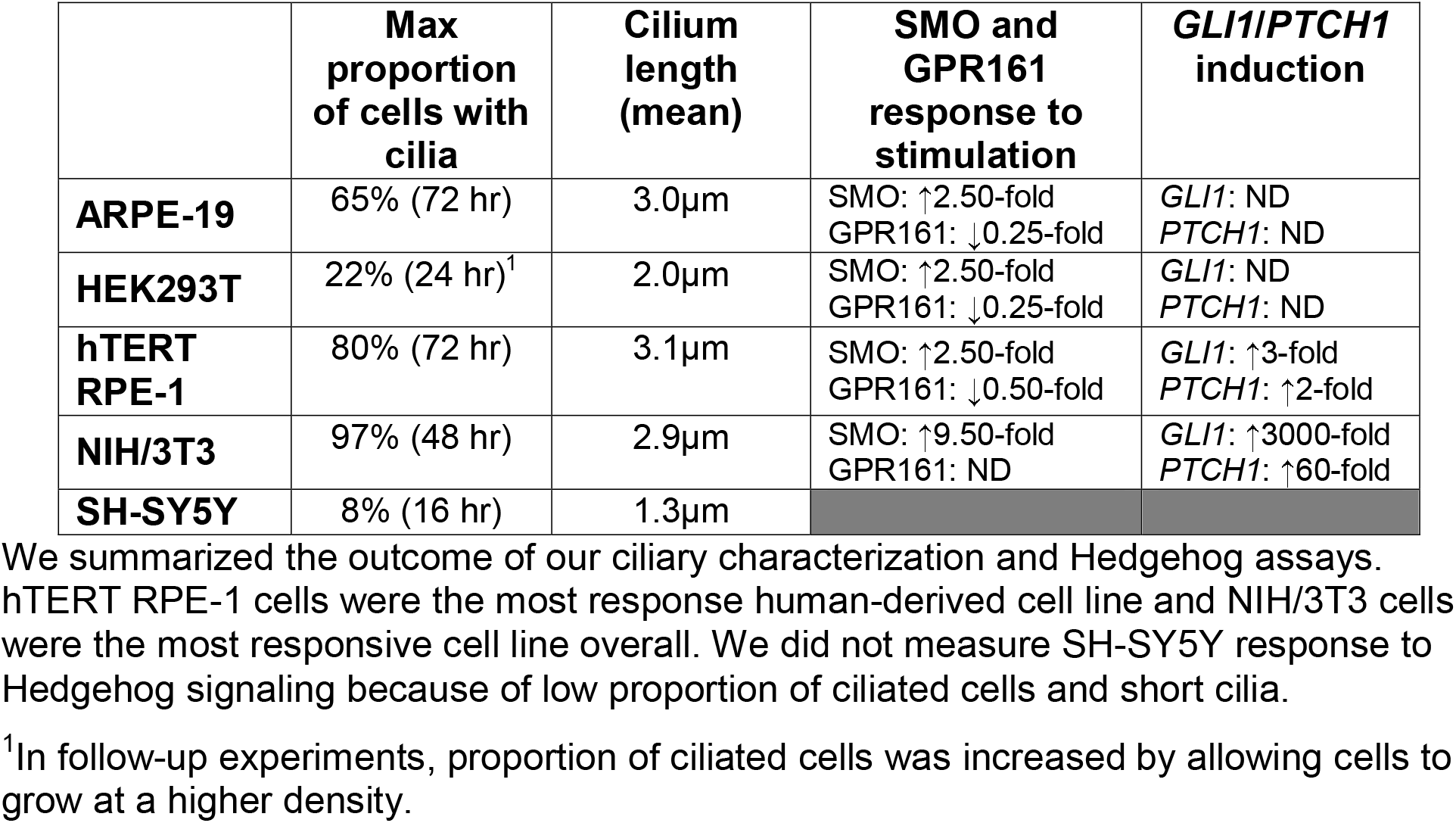
Overview of results.

|  | Max proportion of cells with cilia | Cilium length (mean) | SMO and GPR161 response to stimulation | *GLI1/PTCH1* induction |
| --- | --- | --- | --- | --- |
| **ARPE-19** | 65% (72 hr) | 3.0µm | SMO: ↑2.50-fold<br>GPR161: ↓0.25-fold | *GLI1*: ND<br>*PTCH1*: ND |
| **HEK293T** | 22% (24 hr)<sup>1</sup> | 2.0µm | SMO: ↑2.50-fold<br>GPR161: ↓0.25-fold | *GLI1*: ND<br>*PTCH1*: ND |
| **hTERT RPE-1** | 80% (72 hr) | 3.1µm | SMO: ↑2.50-fold<br>GPR161: ↓0.50-fold | *GLI1*: ↑3-fold<br>*PTCH1*: ↑2-fold |
| **NIH/3T3** | 97% (48 hr) | 2.9µm | SMO: ↑9.50-fold<br>GPR161: ND | *GLI1*: ↑3000-fold<br>*PTCH1*: ↑60-fold |
| **SH-SY5Y** | 8% (16 hr) | 1.3µm |  |  |

We summarized the outcome of our ciliary characterization and Hedgehog assays. hTERT RPE-1 cells were the most responsive human-derived cell line and NIH/3T3 cells were the most responsive cell line overall. We did not measure SH-SY5Y response to Hedgehog signaling because of low proportion of ciliated cells and short cilia.
<sup>1</sup>In follow-up experiments, proportion of ciliated cells was increased by allowing cells to grow at a higher density.

It is also clear from our experiments that each Hh assay requires optimization for each cell line. Prior to assessing Hh response, we had to determine acceptable seeding, growth conditions, and fixation. In HEK293T cells, we had to balance the upside of higher ciliation rates in confluent cultures with the downside of overlapping cells complicating qIF measurements; therefore, low ciliation rates in HEK293T cells may have contributed to the low Hh target gene induction.

For our qIF experiments, we had to identify antibodies that worked across cell lines and were specific to the protein of interest. The limited response to SAG stimulation in ARPE-19 and HEK293T cells could be due to difference in expression of Hh negative regulators in these cell lines that differ from hTERT RPE-1 and NIH/3T3 cells, for example SUFU expression which complexes with GLI2/3 and stabilizes the activator forms of the proteins (28). Alternatively, these cells might respond to native Hh ligand or other agonists, but not SAG. There is prior evidence that hTERT RPE-1 and NIH/3T3 cells relocalize SMO and GPR161 and upregulate pathway target genes in response to Hh stimulation (22,29,30). ARPE-19 cells have been used to determine Hh response to cyclopamine, a down-regulator of the pathway (31). We could not detect Gpr161 in NIH/3T3 cells by immunofluorescence, despite the 94% amino acid identity between the human and mouse for the antigen used to generate the antibody. This may be due to differences between the human and mouse proteins at the key epitopes recognized by the polyclonal antibody. Alternatively, Gpr161 expression has been demonstrated in other studies using different stimulation time points (32). Future work could explore ligand dose-response, and optimal timing for SMO and GPR161 localization in response to Hh stimulation.

qPCR also needs to be tailored to each line. We optimized primer sets, template concentrations, and amplification parameters. Importantly, expression of the commonly used reference gene, *GAPDH*, differed between unstimulated and stimulated HEK293T cells. *GAPDH* expression responds to serum starvation (33,34), adding to the importance of screening reference genes for each cell line. During *GLI1* primer optimization, we identified high CT values in our cell lines (>30 CT), which introduces variability (35). To bring the CT values into a better range, we increased template concentration and added a touchdown step in ARPE-19 and HEK293T, lowering the CT values by 3-4 cycles(17). Unstimulated NIH/3T3 samples also had a CT >30, but the touchdown step did not decrease CT values in this cell line, suggesting the *GLI1* expression is below the level of detection in this cell line and increases substantially with stimulation.

While we confirmed that hTERT RPE-1 and NIH/3T3 cells are appropriate cell lines to model Hh signaling, we have not demonstrated that they recapitulate all aspects of Hh signaling across cell types *in vivo*; therefore, Hh experiments in these cell lines should be validated in additional models, ideally *in vivo*. In addition, we have not fully excluded ARPE-19 and HEK293T cells as potential models, since we did not evaluate all aspects of Hh signaling such as dose-response to ligand, time course, and other important characteristics. If there were compelling reasons to use these cell lines for Hh-related experiments, further characterization might reveal intact Hh responses at different timepoints or different agonist concentrations.

hTERT RPE-1 and NIH/3T3 cells respond consistently to Hh pathway stimulation across two standard assays and are genetically tractable. These cell lines provide models originating from two organisms and could be used individually or in parallel to study Hh response. hTERT RPE-1 cells are a good model system for exploring these mechanistic details at the cellular level.

## Supporting information

**S1 Fig. Extended SH-SY5Y ciliation time course**. Proportion of ciliated cells for SH-SY5Y cells from 0-144 hours. One coverslip was imaged for each timepoint, with number of cells counted at each time point listed below each column in the graph.

**S2 Fig. HEK293T ciliation data**. A) Proportion of ciliated HEK293T cells at 0, 24, 48, and 72 hours post serum starvation from additional thaws. B) Representative images of monolayer areas of HEK293T cell growth from Figure 1 (top row) and repeat experiments with additional thaws of HEK293T cells. Cilia were stained with anti-ARL13B antibody. C) Representative images displaying ciliation in areas with multi-layer growth from additional thaws of HEK293T cells. Cilia were stained with anti-ARL13B antibody. Scale bars are 10μm.

**S3 Fig. GPR161 and SMO localization**. Representative images from each batch showing variation in A) SMO and B) GPR161 localization. C) Normalized fluorescence intensity (arbitrary units, au) of GPR161 with and without stimulation in an additional passage of hTERT RPE-1 cells. Black bars = median normalized fluorescence intensity. Small symbol = normalized fluorescence intensity of one cilium. Numbers of cilia measured are listed below each column in the graph. Representative cilia are shown next to the graphs for unstimulated (-SAG) and stimulated (+SAG) cells. Scale bars are 5 μm. T-test of equal variances was performed. p-values: * 0.01 to 0.05, ** 0.001 to 0.01, and *** <0.001.

**S4 Fig. Variable *GLI1, PTCH1*, and reference gene expression in response to Hh pathway stimulation**. *GLI1, PTCH1*, and reference gene CT (cycle threshold) for ARPE-19, HEK293T, hTERT RPE-1, and NIH/3T3 cell lines with and without stimulation. For cell lines with high CT values (>29), we used a touchdown qPCR method. For each condition (with or without stimulation), we collected mRNA from three batches for each cell line. Normalized *GLI1* and *PTCH1* expression for each qPCR condition is on the far right. Each symbol in the left panels represents the average CT from three technical replicates for each batch. Each symbol in the right panels represents the expression fold change for each batch.

**S1 Data. Experimental data sets**.

## References

1. Berbari NF, O’Connor AK, Haycraft CJ, Yoder BK. The primary cilium as a complex signaling center. Curr Biol CB. 2009 Jul 14;19(13):R526–535.

2. Goetz SC, Anderson KV. The primary cilium: a signalling centre during vertebrate development. Nat Rev Genet. 2010 May;11(5):331–44.

3. Pietrobono S, Franci L, Imperatore F, Zanini C, Stecca B, Chiariello M. MAPK15 Controls Hedgehog Signaling in Medulloblastoma Cells by Regulating Primary Ciliogenesis. Cancers. 2021 Sep 29;13(19):4903.

4. Yoon JW, Gallant M, Lamm MLG, Iannaccone S, Vieux K-F, Proytcheva M, et al. Noncanonical regulation of the Hedgehog mediator GLI1 by c-MYC in Burkitt lymphoma. Mol Cancer Res MCR. 2013 Jun;11(6):604–15.

5. Hulleman JD, Nguyen A, Ramprasad VL, Murugan S, Gupta R, Mahindrakar A, et al. A novel H395R mutation in MKKS/BBS6 causes retinitis pigmentosa and polydactyly without other findings of Bardet-Biedl or McKusick-Kaufman syndrome. Mol Vis. 2016;22:73–81.

6. Breslow DK, Hoogendoorn S, Kopp AR, Morgens DW, Vu BK, Kennedy MC, et al. A CRISPR-based screen for Hedgehog signaling provides insights into ciliary function and ciliopathies. Nat Genet. 2018 Mar;50(3):460–71.

7. Briscoe J, Thérond PP. The mechanisms of Hedgehog signalling and its roles in development and disease. Nat Rev Mol Cell Biol. 2013 Jul;14(7):416–29.

8. Mukhopadhyay S, Wen X, Ratti N, Loktev A, Rangell L, Scales SJ, et al. The Ciliary G-Protein-Coupled Receptor Gpr161 Negatively Regulates the Sonic Hedgehog Pathway via cAMP Signaling. Cell. 2013 Jan 17;152(1):210–23.

9. He M, Subramanian R, Bangs F, Omelchenko T, Liem KF, Kapoor TM, et al. The kinesin-4 protein Kif7 regulates mammalian Hedgehog signalling by organizing the cilium tip compartment. Nat Cell Biol. 2014 Jul;16(7):663–72.

10. Wen X, Lai CK, Evangelista M, Hongo J-A, de Sauvage FJ, Scales SJ. Kinetics of Hedgehog-Dependent Full-Length Gli3 Accumulation in Primary Cilia and Subsequent Degradation. Mol Cell Biol. 2010 Apr 15;30(8):1910–22.

11. Platt KA, Michaud J, Joyner AL. Expression of the mouse Gli and Ptc genes is adjacent to embryonic sources of hedgehog signals suggesting a conservation of pathways between flies and mice. Mech Dev. 1997 Mar;62(2):121–35.

12. Haycraft CJ, Banizs B, Aydin-Son Y, Zhang Q, Michaud EJ, Yoder BK. Gli2 and Gli3 localize to cilia and require the intraflagellar transport protein polaris for processing and function. PLoS Genet. 2005 Oct;1(4):e53.

13. Postma M, Goedhart J. PlotsOfData—A web app for visualizing data together with their summaries. PLOS Biol. 2019 Mar 27;17(3):e3000202.

14. Goedhart J. SuperPlotsOfData—a web app for the transparent display and quantitative comparison of continuous data from different conditions. Pollard T, editor. Mol Biol Cell. 2021 Mar 15;32(6):470–4.

15. Slaats GG, Isabella CR, Kroes HY, Dempsey JC, Gremmels H, Monroe GR, et al. MKS1 regulates ciliary INPP5E levels in Joubert syndrome. J Med Genet. 2016 Jan;53(1):62–72.

16. Schindelin J, Arganda-Carreras I, Frise E, Kaynig V, Longair M, Pietzsch T, et al. Fiji: an open-source platform for biological-image analysis. Nat Methods. 2012 Jun 28;9(7):676–82.

17. Zhang Q, Wang J, Deng F, Yan Z, Xia Y, Wang Z, et al. TqPCR: A Touchdown qPCR Assay with Significantly Improved Detection Sensitivity and Amplification Efficiency of SYBR Green qPCR. PLOS ONE. 2015 Jul 14;10(7):e0132666.

18. Bustin SA, Benes V, Garson JA, Hellemans J, Huggett J, Kubista M, et al. The MIQE guidelines: minimum information for publication of quantitative real-time PCR experiments. Clin Chem. 2009 Apr;55(4):611–22.

19. Jacob F, Guertler R, Naim S, Nixdorf S, Fedier A, Hacker NF, et al. Careful selection of reference genes is required for reliable performance of RT-qPCR in human normal and cancer cell lines. PloS One. 2013;8(3):e59180.

20. Guo S, Liao H, Liu J, Liu J, Tang F, He Z, et al. Resveratrol Activated Sonic Hedgehog Signaling to Enhance Viability of NIH3T3 Cells in Vitro via Regulation of Sirt1. Cell Physiol Biochem Int J Exp Cell Physiol Biochem Pharmacol. 2018;50(4):1346–60.

21. Kilander MBC, Wang C-H, Chang C-H, Nestor JE, Herold K, Tsai J-W, et al. A rare human CEP290 variant disrupts the molecular integrity of the primary cilium and impairs Sonic Hedgehog machinery. Sci Rep. 2018 Nov 26;8(1):17335.

22. Rohatgi R, Milenkovic L, Scott MP. Patched1 Regulates Hedgehog Signaling at the Primary Cilium. Science. 2007 Jul 20;317(5836):372–6.

23. Westlake CJ, Baye LM, Nachury MV, Wright KJ, Ervin KE, Phu L, et al. Primary cilia membrane assembly is initiated by Rab11 and transport protein particle II (TRAPPII) complex-dependent trafficking of Rabin8 to the centrosome. Proc Natl Acad Sci U S A. 2011 Feb 15;108(7):2759–64.

24. Kim S, Zaghloul NA, Bubenshchikova E, Oh EC, Rankin S, Katsanis N, et al. Nde1-mediated inhibition of ciliogenesis affects cell cycle re-entry. Nat Cell Biol. 2011 Apr;13(4):351–60.

25. Takahashi K, Nagai T, Chiba S, Nakayama K, Mizuno K. Glucose deprivation induces primary cilium formation through mTORC1 inactivation. J Cell Sci [Internet]. 018 Jan 8 [cited 2021 Jun 22];131(1). Available from: https://doi.org/10.1242/jcs.208769

26. Corbit KC, Aanstad P, Singla V, Norman AR, Stainier DYR, Reiter JF. Vertebrate Smoothened functions at the primary cilium. Nature. 2005 Oct 13;437(7061):1018– 21.

27. Livak KJ, Schmittgen TD. Analysis of relative gene expression data using real-time quantitative PCR and the 2(-Delta Delta C(T)) Method. Methods San Diego Calif. 2001 Dec;25(4):402–8.

28. Wang C, Pan Y, Wang B. Suppressor of fused and Spop regulate the stability, processing and function of Gli2 and Gli3 full-length activators but not their repressors. Dev Camb Engl. 2010 Jun;137(12):2001–9.

29. Frasca A, Spiombi E, Palmieri M, Albizzati E, Valente MM, Bergo A, et al. MECP2 mutations affect ciliogenesis: a novel perspective for Rett syndrome and related disorders. EMBO Mol Med. 2020 Jun 8;12(6):e10270.

30. Hey CAB, Larsen LJ, Tümer Z, Brøndum-Nielsen K, Grønskov K, Hjortshøj TD, et al. BBS Proteins Affect Ciliogenesis and Are Essential for Hedgehog Signaling, but Not for Formation of iPSC-Derived RPE-65 Expressing RPE-Like Cells. Int J Mol Sci. 2021 Jan 29;22(3):1345.

31. Duan F, Lin M, Li C, Ding X, Qian G, Zhang H, et al. Effects of inhibition of hedgehog signaling on cell growth and migration of uveal melanoma cells. Cancer Biol Ther. 2014 May;15(5):544–59.

32. Hong S-R, Wang C-L, Huang Y-S, Chang Y-C, Chang Y-C, Pusapati GV, et al. Spatiotemporal manipulation of ciliary glutamylation reveals its roles in intraciliary trafficking and Hedgehog signaling. Nat Commun. 2018 Apr 30;9(1):1732.

33. Schmittgen TD, Zakrajsek BA. Effect of experimental treatment on housekeeping gene expression: validation by real-time, quantitative RT-PCR. J Biochem Biophys Methods. 2000 Nov 20;46(1–2):69–81.

34. Krzystek-Korpacka M, Hotowy K, Czapinska E, Podkowik M, Bania J, Gamian A, et al. Serum availability affects expression of common house-keeping genes in colon adenocarcinoma cell lines: implications for quantitative real-time PCR studies. Cytotechnology. 2016 Dec;68(6):2503–17.

35. Taylor SC, Nadeau K, Abbasi M, Lachance C, Nguyen M, Fenrich J. The Ultimate qPCR Experiment: Producing Publication Quality, Reproducible Data the First Time. Trends Biotechnol. 2019 Jul;37(7):761–74.

