## Supplementary figures and images for "Systematic analysis of cilia characteristics and Hedgehog signaling in five immortal cell lines"

### Supplementary Figure 1

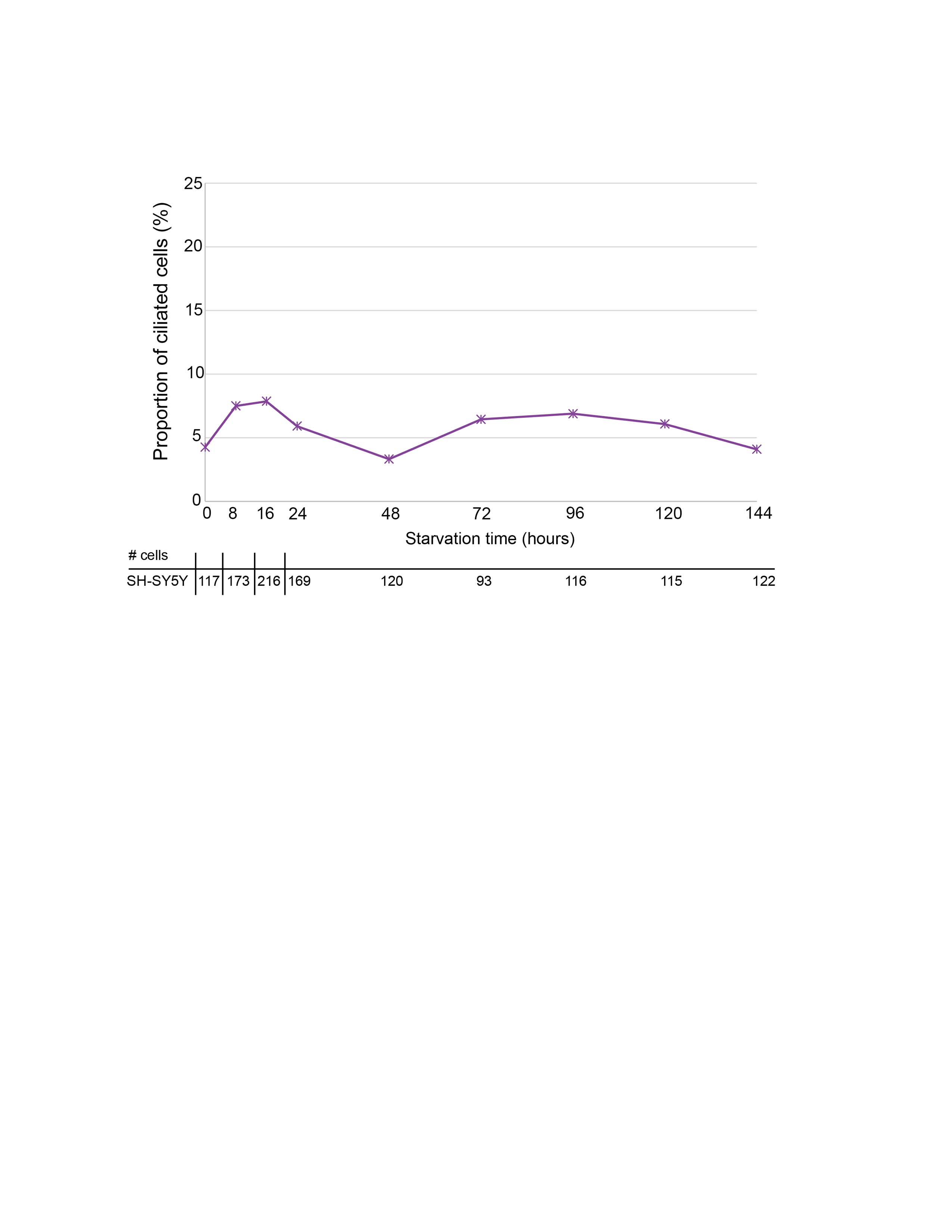

### Supplementary Figure 2

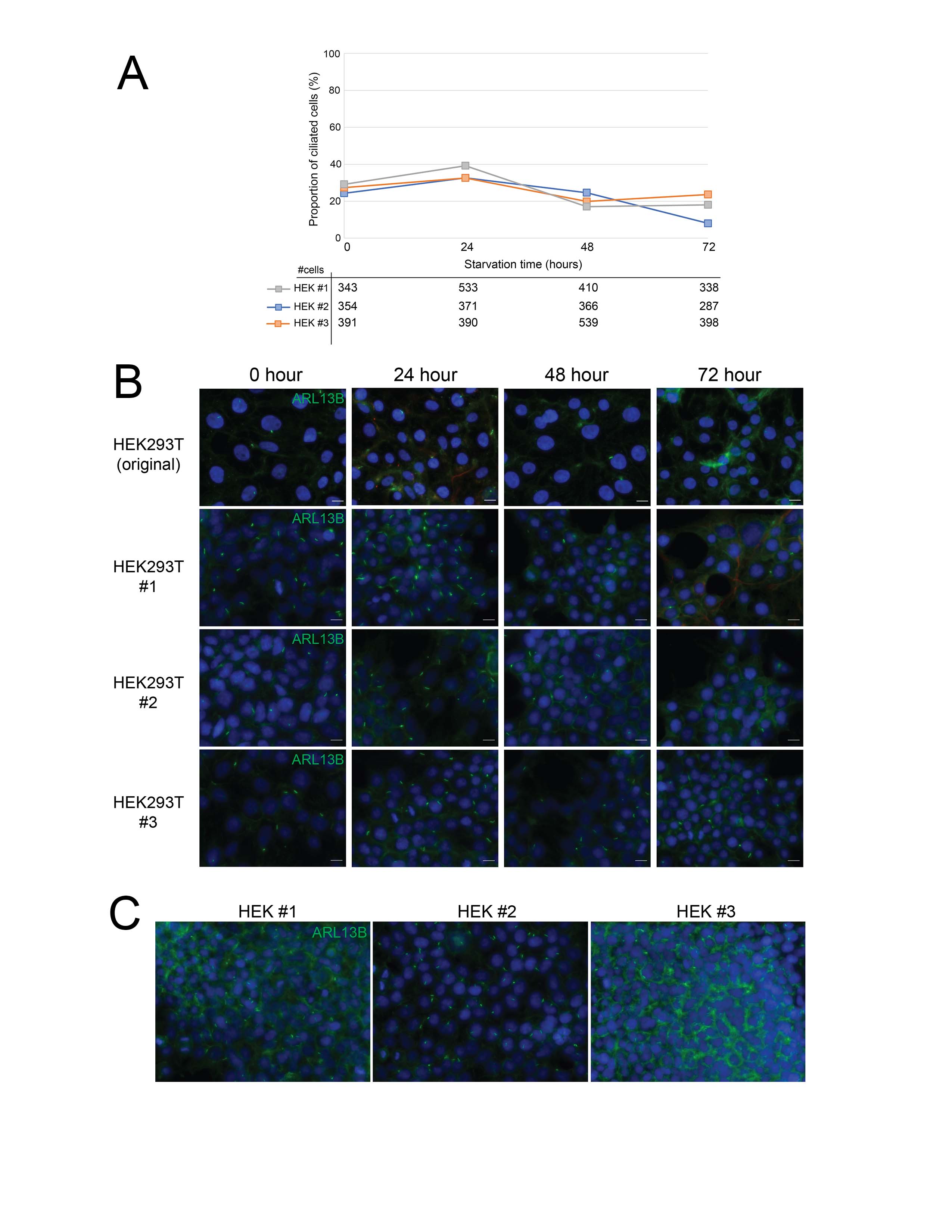

### Supplementary Figure 3

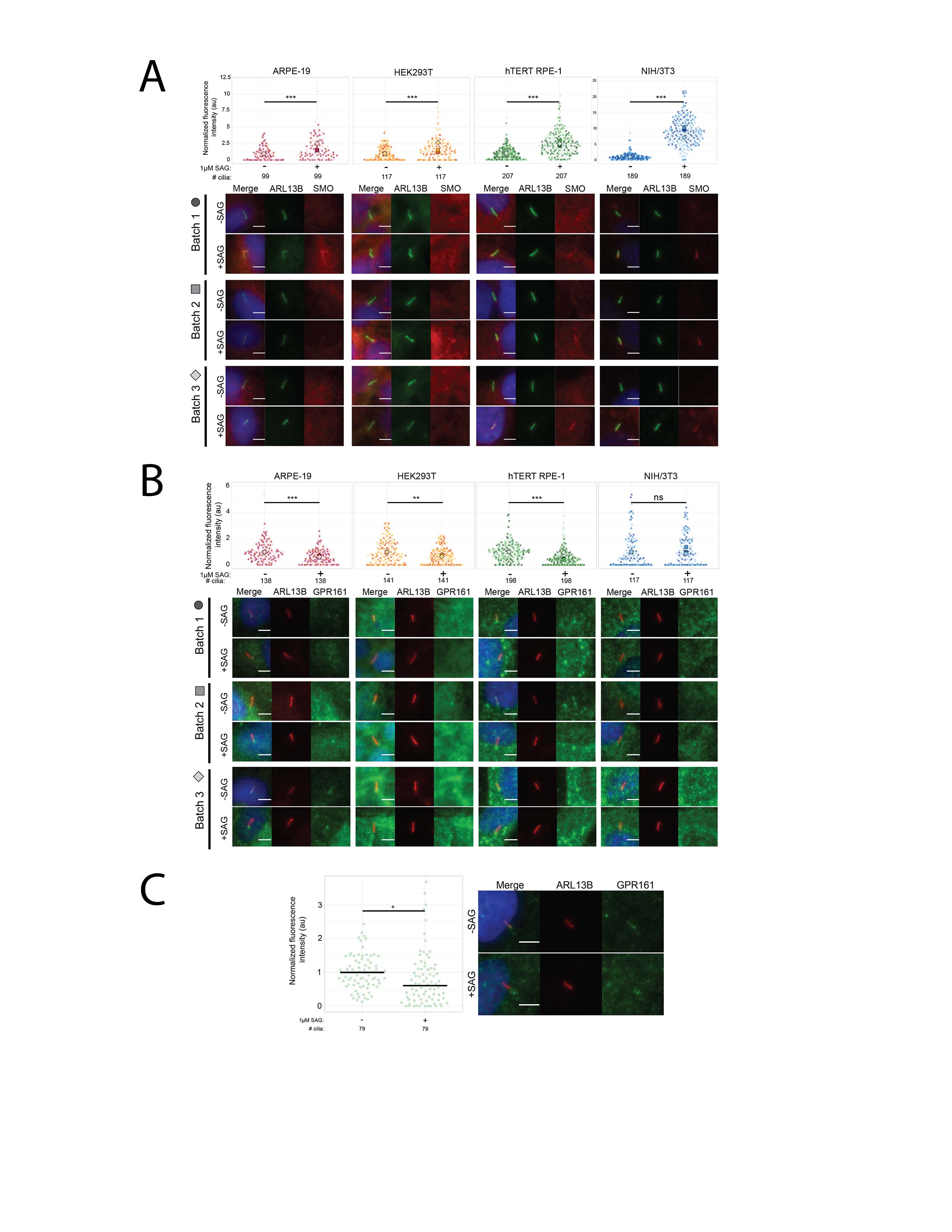

### Supplementary Figure 4

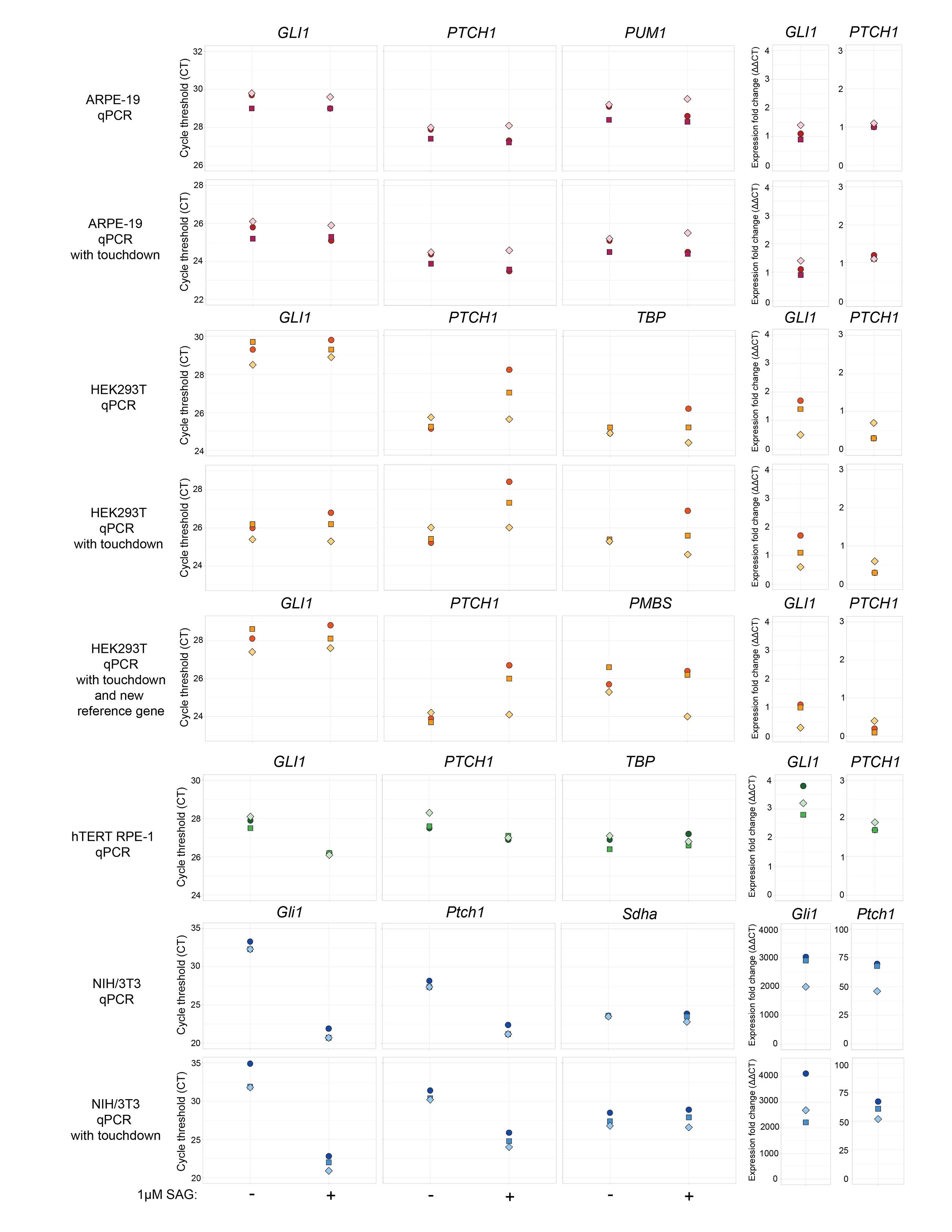
